# Rectal Transplantation of Fragmented Intestinal Organoids and Crypts Promotes Mucosal Regeneration in DSS-Induced Colitis

**DOI:** 10.1101/2025.11.15.688663

**Authors:** Yihan Wang, Xuesen Xu, Dong Wang

## Abstract

**Background:** Ulcerative colitis (UC) represents a global health challenge characterized by relapsing inflammation and epithelial barrier disruption. Conventional immunosuppressive therapies have reached their therapeutic ceiling, with mucosal healing and long-term remission remaining unmet therapeutic goals for many patients. Organoid-based regenerative therapy offers a new paradigm by reconstructing damaged mucosa rather than merely controlling inflammation.

**Methods:** We hypothesized that fragmented intestinal organoids, building upon the foundational model established by Yui et al., whose use of immunodeficient Rag2^−^/^−^ mice and surgical mucosal injury first demonstrated the feasibility of colon organoid transplantation. Our study advances this framework by introducing optimized transplantation timing and employing immunocompetent C57BL/6 mice to refine translational relevance and clinical feasibility, thereby extending the applicability of organoid therapy to more realistic physiological contexts., when transplanted rectally at an optimized time point, could integrate more efficiently into inflamed mucosa and accelerate epithelial regeneration even in an immunocompetent environment. A 2.5% dextran sulfate sodium (DSS)-induced colitis model was established in C57BL/6 mice. Intestinal organoids and crypts derived from EGFP-transgenic donors were cultured and transplanted rectally using a minimally invasive delivery system. Fluorescence microscopy, histological analysis, and clinical indices were used to evaluate engraftment and mucosal repair.

**Results:** EGFP^+^ organoid fragments and crypts successfully engrafted at ulcerated sites, reconstituting epithelial structures and restoring mucosal integrity. Mice receiving organoid or crypts transplantation exhibited rapid weight recovery and reduced bleeding compared to DSS-only controls. Histological and fluorescence analyses confirmed epithelial restitution exceeding 60% by day 7 post-administration versus 18% in controls. These results validate the regenerative potential of organoid transplantation within an immunocompetent host.

**Conclusion:** This study demonstrates that rectal transplantation of fragmented intestinal organoids and crypts promotes robust mucosal regeneration in DSS-induced colitis. By optimizing transplantation timing and employing a clinically relevant model, our work establishes a translational foundation for organoid-based regenerative therapies targeting inflammatory bowel diseases.

## Introduction

Inflammatory bowel diseases (IBDs) is a chronic condition that causes relapsing inflammation in gastrointestinal tract. Crohn’s disease (CD) and ulcerative colitis (UC) are the two main forms of IBD^1^. The prevalence of IBD varies between countries and has been constantly increasing in the world. By 2025, prevalence in North America will exceed 0.8%,while incidence in newly industrialized regions continues to climb under the pressures of urbanization and dietary westernization. Combined with the aging demographic of the IBD patient population, this epidemiological shift suggests that by 2045, IBD will remain a sustained global burden^2^. Chronic inflammatory processes of the intestinal tract of unknown origin characterize both CD and UC. IBD is known for its recurrent flare-ups and can be associated with chronic diarrhea, abdominal pain, rectal bleeding, fatigue, and extraintestinal manifestations, such as arthritis, uveitis, and skin lesions, severely impacting patients’ quality of life^3^. Although molecular insights and biologics have advanced treatment outcomes, current immunosuppressive therapies have plateaued: for many patients, mucosal healing and durable remission remain out of reach, and surgery often becomes the last resort^4^.

Given these limitations, regenerative medicine offers a new frontier. Organoids are a group of stem cell-originated 3D cell clusters, which maintain the functionality and heterogeneity of the originating organs, intestinal organoids can be generated from adult stem cells or pluripotent stem cells ^5^. Furthermore, intestinal organoids derived from pluripotent stem cell, either embryonic stem cells or induced pluripotent stem cells, contain also epithelial cells and mesenchymal cells, whereas adult stem cells formed intestinal organoids are only epithelial cell composed ^6^. Intestinal organoids are thus cell-heterogeneous, allowing for wide application in host-microbiome interaction research, gene editing, molecular signaling analysis and immunological research^7^. Recently, study has reported the use of intestinal organoid to ameliorate mouse colitis, which transplant intestinal organoids into a mouse model of colitis ^8^.Foundational progress was made by Yui et al. in a landmark 2012 Nature Medicine study, where the TMDU and Hubrecht Institute team first achieved long-term expansion of Lgr5^+^ colonic stem cells and demonstrated functional engraftment of cultured colon organoids into damaged murine colon^9^. Their work provided proof-of-concept that adult intestinal stem cells could reconstruct the epithelial barrier in vivo, laying the groundwork for subsequent translational advances in organoid-based regenerative therapy. Intestinal organoid transplantation, leveraging the self-renewal and multilineage differentiation of intestinal stem cells, provides an opportunity to reconstruct epithelial architecture and restore barrier function. Yet, three persistent barriers hinder clinical translation: determining the optimal transplantation window, ensuring survival within inflamed tissue, and validating efficacy in immunocompetent hosts.

We hypothesized that fragmented intestinal organoids, delivered rectally at an optimized window, would integrate efficiently into inflamed mucosa and promote rapid epithelial regeneration. This study introduces two methodological innovations that address prior limitations. First, we used daily morphological assessment under fluorescence microscopy to non-destructively identify the ideal transplantation timing, maximizing regenerative potential. Second, unlike previous studies that relied on immunodeficient mice, we used immunocompetent C57BL/6 mice,including both male and female donors,to more accurately minic clinical conditions.

## Materials and Methods

### Animals

B6-H11-CAG-EGFP (donor) and C57BL/6J (recipient) mice were purchased from the GemPharmatech company and maintained under SPF conditions. All procedures were approved by the Animal Ethics Committee of China Pharmaceutical University.

### Organoid culture

Intestinal crypts were isolated from EGFP^+^ donor mice, embedded in Matrigel, and cultured in complete IntestiCult™ medium (Stem Cell, #06005). Organoids were maintained by routine passaging every 6–7 days to ensure long-term expansion and stability. On day 6 of each culture cycle, organoids exhibiting vigorous budding were divided into two fractions: one fraction was used immediately for transplantation experiments, while the remaining fraction was mechanically fragmented and re-plated to initiate the next passage. This repeated expansion maintained robust viability and proliferative capacity across multiple passages, demonstrating that donor-derived organoids preserved their regenerative potential throughout extended culture.

### Colitis induction

Acute colitis was induced by providing 2.5% DSS (w/v) in drinking water for 7 days. Control mice received regular water.Mice were monitored daily, and transplantation was performed once body weight loss reached ≥10%, indicating successful induction of acute colitis.

### Rectal transplantation

Under avertin anesthesia, 100 μL of organoid suspension (≈1–3×10^6^ cells/mouse) was delivered rectally via a lubricated 300 μm catheter (∼4 cm depth). The anus was sealed with a thin layer of medical scar gel for 24 hours to prevent leakage.

### Evaluation

Body weight and stool consistency were monitored and documented daily. On day 7 post-transplantation, mice were euthanized, and colon length, histology (H&E), and GFP fluorescence were analyzed.

## Results

### Comparison of colonic and ileal organoids

To establish a consistent and high-quality donor source for organoid generation, intestinal crypts were isolated from freshly dissected small intestine and colon tissues following a modified protocol adapted from the *Nature Protocols*paper by Sato et al.^8^. Briefly, the intestinal lumen was flushed with cold PBS, opened longitudinally, and cut into small pieces (2–4 mm). Tissue fragments were washed repeatedly with cold PBS until the supernatant became clear to remove debris and villi. Crypts were released by gentle shaking in chelation buffer containing 2 mM EDTA, filtered through a 70 μm cell strainer, and pelleted by centrifugation at 300 × g for 5 min. The isolated crypts were then embedded in Matrigel and cultured in IntestiCult™ medium at 37°C with 5% CO_2_.

To determine the most suitable organoid source for transplantation, we first compared the growth characteristics of colonic and ileal organoids cultured under identical conditions. Morphological assessment revealed that ileal organoids exhibited higher initial plating density and faster reformation after passage compared with colonic organoids, which appeared more cystic and fragile. By day 3, ileal organoids displayed greater structural integrity and budding frequency, whereas colonic organoids remained predominantly spherical with limited branching (Figure 1a). Despite ulcerative colitis being a colonic disease, these observations led us to select ileal organoids as the donor source for subsequent experiments due to their superior growth vigor and stability.

**Figure 1.**
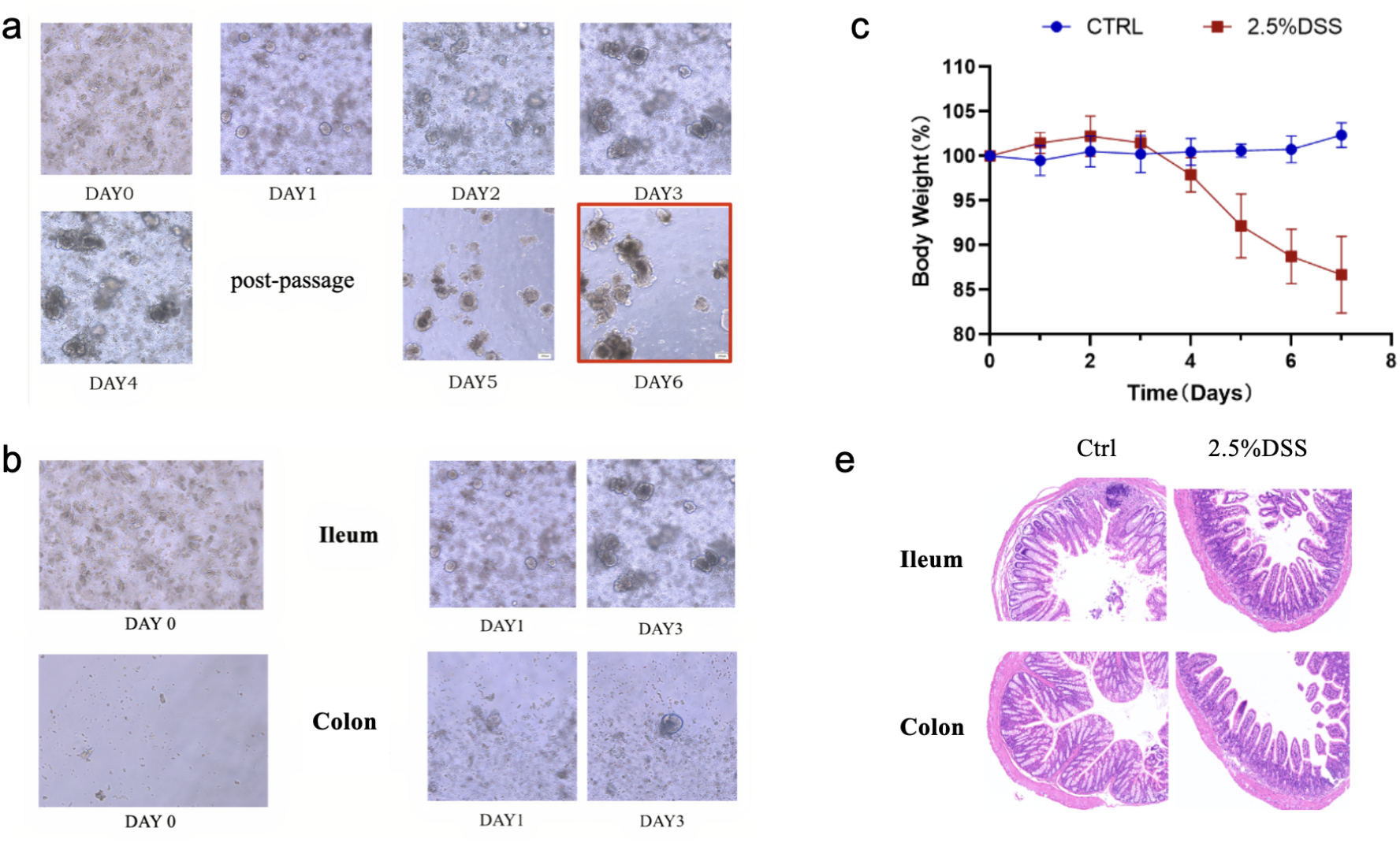
Morphological comparison and colitis model validation. (a) Time-course morphological evolution of intestinal organoids post-passage (Day 0–6), showing progressive cyst formation, budding, and polarization, with Day 6 identified as the optimal transplantation window. (b) Comparative morphology of colonic versus ileal organoids during early culture (Day 0–3), illustrating faster proliferation and denser growth in ileal organoids. (c) Validation of DSS-induced colitis model by monitoring body weight loss in mice receiving 2.5% DSS compared with control group. (e) Representative H&E staining of colon and ileum sections from control and DSS-treated mice demonstrating epithelial disruption, crypt distortion, and inflammatory infiltration in DSS-induced colitis compared with normal tissue architecture.

### Optimized transplantation window

Using morphological evaluation rather than destructive assays, we identified day 6 of organoid culture as the optimal transplantation point, corresponding to maximal budding complexity and epithelial polarity. Daily imaging after passage revealed a distinct morphogenetic trajectory. On day 0, isolated crypt fragments appeared as irregular aggregates with limited structural definition. By day 1–2, spherical cyst-like structures began to emerge, displaying early lumen formation and uniform epithelial borders. On day 3, multiple buds protruded from the cysts, indicating the onset of crypt-like domain formation(Figure 1a). From day 4 to day 5, these organoids expanded rapidly with increased optical density(surface area) and complex branching(budding), demonstrating vigorous proliferative and differentiative activity. By day 6, the organoids exhibited highly defined budding architecture with visible apical–basal polarity, dense epithelial cell layering, and robust three-dimensional organization, which features indicative of peak regenerative potential(Figure 1b). This timing preserved proliferative vigor while ensuring differentiation readiness, serving as the critical window for transplantation to maximize engraftment success^8,9^.

### Validation of DSS-induced colitis

DSS–induced colitis is a well-established murine model that closely mimics human ulcerative colitis by disrupting the epithelial barrier and triggering innate immune activation in the colon. Oral administration of DSS in drinking water results in epithelial erosion, ulceration, and infiltration of inflammatory cells, leading to body weight loss, rectal bleeding, and shortened colon length. This chemically induced model provides a reproducible and controllable system to study mucosal injury and repair mechanisms in vivo^10^.

To verify the establishment of the ulcerative colitis model, mice were divided into four groups according to treatment protocol (Table 1). The control group received no treatment. The DSS group was administered 2.5% DSS solution from Day 0 to Day 7 without rectal transplantation. The crypt group underwent the same DSS induction followed by rectal delivery of crypt fragments isolated from the small intestine of EGFP-transgenic mice on Day 7. The organoid group received DSS induction and rectal transplantation of intestinal organoid fragments on Day 7(Figure 2).

**Table 1.** Experimental groups and treatment protocols. Mice were divided into four groups for evaluation of DSS-induced colitis and organoid transplantation outcomes. The control group received no treatment. The DSS group underwent DSS induction without rectal infusion. The crypt group received rectal administration of crypt fragments from EGFP-transgenic mice after DSS induction. The organoid group received rectal transplantation of intestinal organoid fragments derived from EGFP-transgenic mice on Day 7 following DSS induction.

| Group | Treatment description |
| --- | --- |
| Ctrl group | No treatment was performed. |
| DSS group | DSS-induced colitis was performed from Day 0 to Day 7 without rectal administration. |
| Crypt group | DSS-induced colitis was performed from Day 0 to Day 7; on Day 7, rectal administration of crypt fragments isolated from the small intestine of EGFP-transgenic mice was performed. |
| **Organoid group** | DSS-induced colitis was performed from Day 0 to Day 7; on Day 7, rectal administration of intestinal organoid fragments derived from EGFP-transgenic mice was performed. |

**Figure 2.**
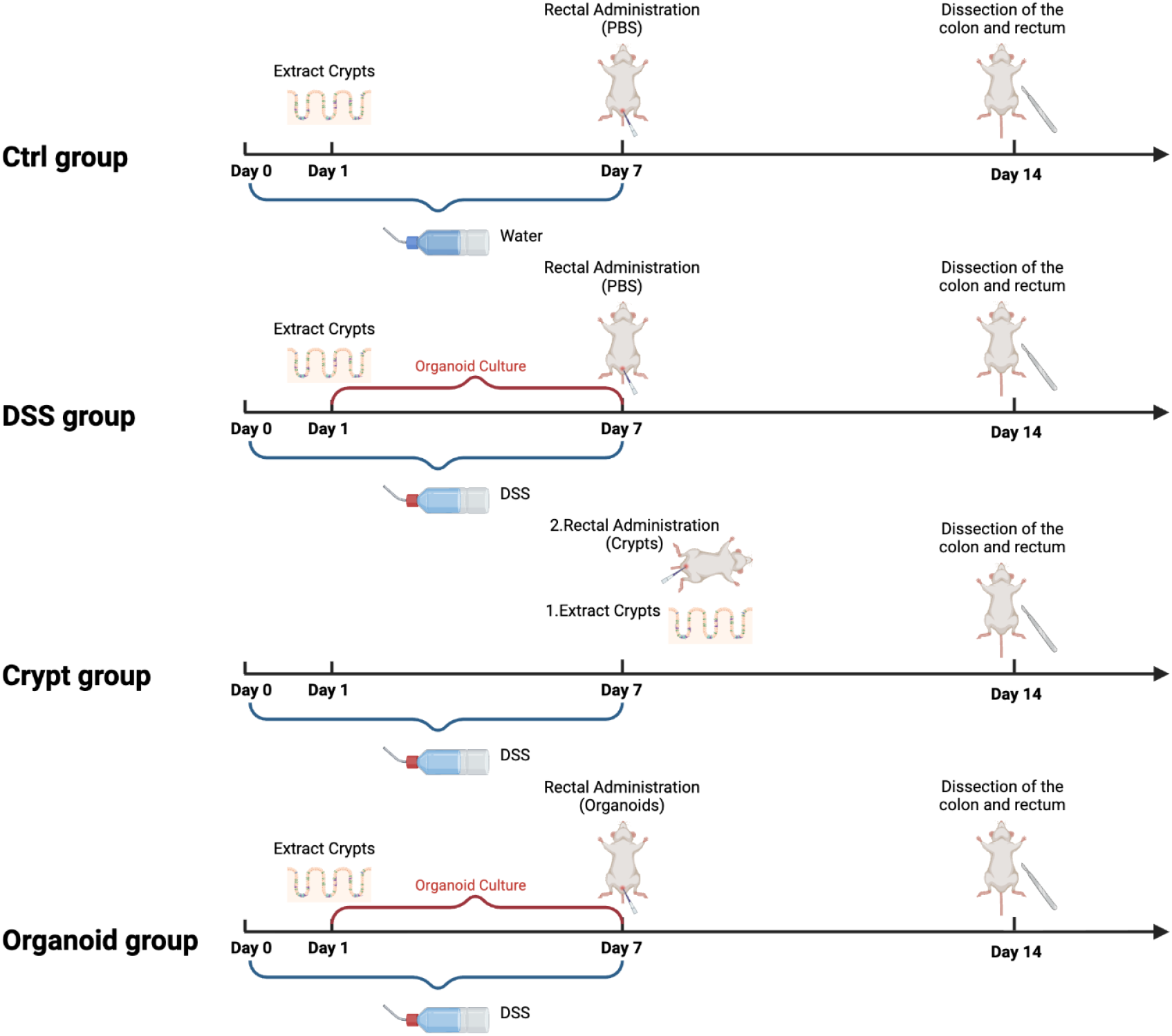
Overview of experimental workflow and group-specific interventions in the DSS-induced colitis model

C57BL/6J mice exposed to DSS developed hallmark features of UC including body weight loss (∼15%), rectal bleeding, and shortened colon length—confirming successful disease induction (Figure 1c).H&E staining of colon and ileum sections further verified the pathological features of DSS-induced colitis, revealing epithelial erosion, crypt atrophy, and inflammatory cell infiltration compared with intact mucosa in control mice (Figure 1e)^11^.

### Regenerative outcomes following organoid transplantation

Following rectal transplantation, mice that received crypts fragments displayed significant clinical improvement compared with the DSS-only group. Body weight recovery was observed within 3–5 days after transplantation, and stool consistency progressively normalized (Figure 3a). Both the crypt and organoid groups exhibited an upward trend in body weight after Day 11, indicating gradual recovery, while the DSS-only group showed only transient improvement followed by continuous decline. This suggests that transplantation effectively mitigated disease progression and promoted systemic recovery. Macroscopic examination of colons revealed visibly reduced inflammation and greater tissue integrity in organoid-treated mice, with colon lengths notably preserved compared with DSS controls (Figure 3b–c). These gross anatomical differences were obtained from separate experimental batches, reflecting consistent regenerative effects across replicates. Collectively, these findings indicate that organoid transplantation alleviates disease severity and promotes recovery of mucosal structure and function.Gross inspection of the colon revealed visibly less edema, ulceration, and bleeding in organoid-treated mice, accompanied by a notable preservation of colon length compared with DSS controls (Figure 3b–c). Both the crypt and organoid groups exhibited colon and rectum lengths approaching those of the control group, indicating substantial anatomical recovery and reduced inflammatory contraction of the intestinal tissue. These macroscopic findings, collected from separate experimental batches, consistently indicated that organoid transplantation alleviated disease severity, reduced inflammatory damage, and promoted overall recovery of the intestinal structure and function.

**Figure 3.**
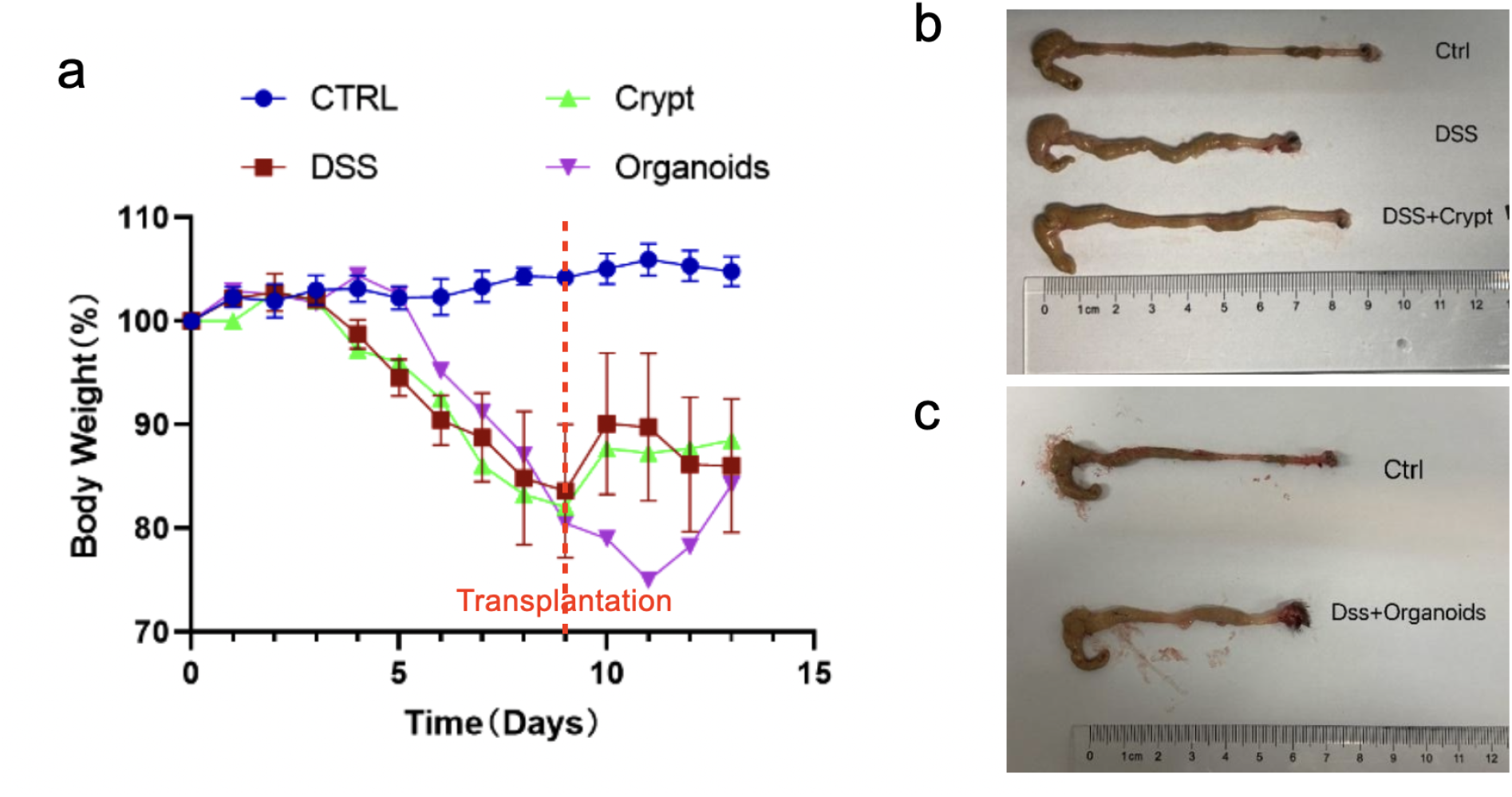
Clinical and anatomical recovery following organoid transplantation. (a) Body weight trajectory of mice from the four experimental groups (Ctrl, DSS, Crypt, Organoid) with the transplantation point indicated by a dashed red line. Both the crypt and organoid groups displayed steady post-transplantation weight recovery, whereas the DSS group showed only transient improvement followed by decline. (b–c) Representative macroscopic images of colons collected from two independent experimental batches showing that both the crypt and organoid groups exhibited colon and rectum lengths close to those of control mice, indicating substantial anatomical recovery and reduced inflammatory contraction of intestinal tissue.

### Epithelial integration and mucosal reconstruction

Before transplantation, EGFP^+^ donor crypts and organoids retained robust fluorescence for at least five days in culture, confirming stable reporter expression for subsequent in vivo tracking (Figure 4a). EGFP fluorescence tracking revealed that donor-derived epithelial cells successfully integrated into the damaged mucosal surface, forming continuous epithelial layers that bridged ulcerated regions (Figure 4b). Fluorescent signal localization along crypt-villus domains indicated that transplanted cells actively participated in barrier reconstruction rather than remaining as isolated grafts. Corresponding H&E staining further confirmed organized epithelial regeneration, reduced inflammatory infiltration, and re-established crypt architecture (Figure 3b). Collectively, these results demonstrate that rectally delivered organoid fragments not only survive within the inflammatory milieu but also integrate functionally into host tissue to achieve mucosal restoration.

**Figure 4.**
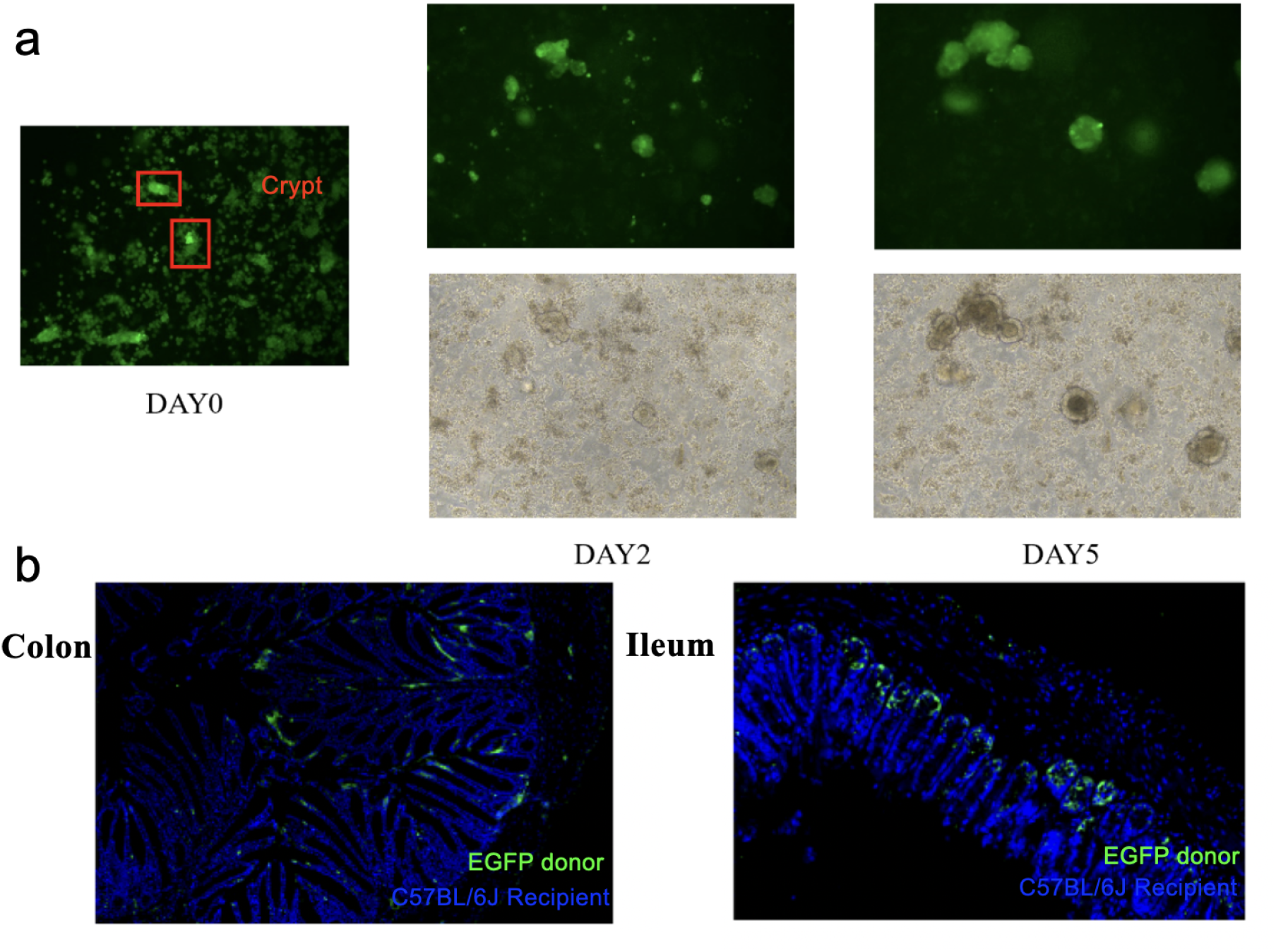
Epithelial integration and mucosal reconstruction. (a) Fluorescence and bright-field images of EGFP-derived crypts and their derived organoids at Day 0, Day 2, and Day 5. Freshly isolated crypts display strong green fluorescence under fluorescence microscopy at Day 0, and the signal remains readily detectable through Days 2 and 5 during culture, indicating stable and persistent reporter expression in donor tissue used for transplantation. (b) Representative fluorescence micrographs of colon and ileum tissues from recipient mice after rectal transplantation, showing green EGFP^+^ donor-derived epithelial cells integrated within host mucosa. Reconstructed epithelial layers align along the crypt–villus axis, confirming functional engraftment and mucosal repair in DSS-induced colitis.

## Discussion and conclusion

Our study demonstrates that fragmented ileal organoids and isolated crypts, transplanted rectally at an optimized timing, can regenerate the damaged mucosa in an immunocompetent model of ulcerative colitis. This approach addresses key translational barriers in organoid-based therapy. Three technical innovations define our work. First, the non-destructive morphological assessment enables real-time evaluation of organoid readiness for transplantation, offering a simple yet precise optimization strategy. Second, the use of immunocompetent mice introduces an immune context that mirrors human physiology, enhancing clinical relevance. Third, we selected ileal organoids as the donor source based on their superior proliferative vigor, structural stability, and robust repair capacity. Despite being derived from the small intestine, ileal organoids are fully capable of integrating into and repairing colonic lesions, and critically they expand more readily than colonic organoids, making them more suitable for large-scale production and translational application^12^. Together, these refinements advance organoid transplantation toward translational feasibility.

The observed epithelial regeneration likely reflects activation of endogenous repair pathways, including Wnt/EGF signaling and epithelial cell migration. The consistent improvement in clinical symptoms, body weight recovery, and macroscopic healing indicates that transplanted organoid fragments facilitated both structural and functional repair of the intestinal barrier. The crypt and organoid groups exhibited nearly restored colon and rectum lengths comparable to control mice, underscoring effective anatomical recovery and reduced inflammatory contraction^13^. These results collectively highlight the efficacy of organoid-based transplantation in mitigating disease severity and enhancing mucosal restoration. Even in the absence of fluorescent tracking, the preservation of tissue integrity and reduction of inflammatory infiltration substantiate that regeneration was driven by successful epithelial restitution rather than transient paracrine effects.

We also included a crypt transplantation group in our design to serve as a biologically relevant comparison for organoid therapy. The crypt fragments, composed primarily of freshly isolated intestinal stem cells and progenitors, provide a simpler yet less mature cellular composition than organoids. By including this group, we sought to distinguish the regenerative capacity of unexpanded crypts from that of fully developed organoid fragments, thereby evaluating whether in vitro maturation enhances mucosal repair efficacy. However, due to time constraints, we were unable to conduct a detailed comparative analysis between the two transplantation strategies. Interestingly, preliminary observations, such as body weight recovery, indicated that the crypt group exhibited slightly better outcomes than the organoid group.These findings prompt the question of whether freshly isolated crypts may contain transient pro-regenerative signaling factors or cell populations, potentially paracrine cues supporting epithelial attachment and repair,that are diminished or lost during in vitro organoid maturation. Although our study was not designed to dissect these mechanisms, the trend suggests a biologically meaningful difference that warrants dedicated investigation in future work.

Additionally, our study incorporated both sexes of donor mice to ensure reproducibility and biological diversity. Female EGFP-transgenic mice served as donors for the crypt transplantation group, while male C57BL/6 mice were used as donors for the organoid transplantation group. This design minimized sex bias and provided complementary validation across independent donor sources, further supporting the robustness of our conclusions.

While our findings confirm robust acute-phase regeneration, further studies are needed to evaluate long-term integration, lineage stability, and immune adaptation. Chronic colitis models and direct comparison with pharmacological standards (e.g., anti-TNF agents) will help position organoid transplantation within the therapeutic landscape. In the future, identifying clinically compatible and immunologically appropriate cell sources will be essential. Autologous organoids offer immune matching but may have reduced proliferative capacity in patients with severe or chronic disease. iPSC(induced pluripotent stem cells)-derived intestinal organoids provide a theoretically unlimited and patient-specific source, though their maturation state and functional fidelity require further optimization^14^. Gene-edited allogeneic donor organoids may also reduce immunogenicity and support the development of standardized, banked grafts^15^.

An additional translational challenge lies in the culture system itself. Current intestinal organoid expansion relies heavily on Matrigel, a mouse sarcoma–derived extracellular matrix that is compositionally undefined and unsuitable for clinical use due to batch variability and xenogeneic origin. As highlighted in recent work, transitioning toward chemically defined, xeno-free, and regulatory-compliant matrices, such as synthetic hydrogels or recombinant ECM substitutes, will be crucial for scaling organoid production and enabling clinical-grade manufacturing^16^.

Taken together, rectal transplantation of fragmented intestinal organoids derived from EGFP donor mice effectively repaired DSS-induced mucosal injury. By optimizing transplantation timing and modeling in immunocompetent hosts, this strategy demonstrates a minimally invasive and translationally relevant approach for mucosal regeneration in IBD.

### Patent

This work has been filed under Chinese Patent Application No. CN 119548529.

